# MeFluHyA: a novel fluorescent screening tool for high-throughput identification of HDAC6-selective inhibitors

**DOI:** 10.64898/2026.07.24.740064

**Authors:** Yvonne E. Klingl, Jozefien Goethals, Adrià Sicart, Carlotta Borgarelli, Joris Van Lindt, Robert Prior, Ermal Ismalaj, Wim M. De Borggraeve, Philip Van Damme, Jacob M. Hooker, Michele Curcio, Steven Verhelst, Matthias Schoenberger, Ludo Van Den Bosch

## Abstract

The cytosolic histone deacetylase 6 (HDAC6) plays a key role not only in cancer but also in neurodegenerative diseases, including amyotrophic lateral sclerosis (ALS) peripheral neuropathies. Pharmacological inhibition as well as genetic silencing of HDAC6 is able to rescue several defects. Therefore, developing pharmacological compounds targeting this enzyme is of crucial interest. Here, we report the design, synthesis, and characterization of Methyl Fluorescent Hydroxamic Acid (MeFluHyA), a novel fluorescent HDAC6-selective probe designed for cell imaging. By integrating a Cy5 fluorophore into a phenyl hydroxamic acid scaffold, MeFluHyA shows binding to the catalytically active CD2 domain of HDAC6 without significantly inhibiting its deacetylating function in the sub-micromolar range. Biochemical enzymatic activity assays confirmed its selectivity over other HDAC isoforms. Fluorescent imaging studies in HeLa cells demonstrated strong colocalization with HDAC6-eGFP and a commercial HDAC6 antibody. Competitive binding assays revealed that MeFluHyA effectively identifies known HDAC6 inhibitors, with reduced probe-binding serving as a readout for successful target engagement. MeFluHyA is a cell-permeable, fast and easy to use probe that can be added organism-independent, and its red-shifted fluorophore makes it compatible with multiplex, live and fixed imaging. Overall, we provide a novel tool for screening platforms aimed at identifying novel HDAC6-targeting inhibitors.

## Introduction

Histone deacetylases (HDACs) are a family of enzymes mainly responsible for the deacetylation of histones. Although most HDACs are critical regulators of transcription, HDAC6 stands out structurally and functionally. First, HDAC6 has not one, but two independent catalytic deacetylating domains (CD1 and CD2) with a substrate-specific activity^1,2^. Second, HDAC6 contains a zinc-finger ubiquitin-binding domain (ZnFUBP), involved in protein clearance and degradation via binding of misfolded proteins. Within the two deacetylating domains, a small dynein motor binding domain (DMB) is located, binding dynein and providing transport of misfolded ubiquitinated proteins to aggressomes^3^. Third, the enzyme has two nuclear export signals (NES) and a cytoplasmic retention signal (SE14), responsible for the protein’s cytoplasmic subcellular localization^2^. Interestingly, due to its subcellular localization, HDAC6 deacetylates a variety of non-histone substrates, including cytoskeletal proteins such as α-tubulin, tau and cortactin, as well as proteins involved in protein degradation, such as the chaperone heat shock protein 90 (Hsp90) and redox regulators such as peroxiredoxins^1,2,4–10^.

HDAC6 has been implicated in different neurodegenerative diseases such as Alzheimer’s, Parkinson’s and Huntington’s disease, amyotrophic lateral sclerosis (ALS) and Charcot-Marie-Tooth disease (CMT)^2,3,11–19^. Pharmacological inhibition as well as genetic silencing of HDAC6 has a therapeutic effect in mouse models of inherited neuropathies and chemotherapy-induced peripheral neuropathies^20^. In addition, inhibition of HDAC6 rescues the axonal transport defects as well as other phenotypes observed in various patient-derived ALS models^2,14–16,21–25^. Also in other diseases, such as cancer^26^, heart failure^27^, ischemia/reperfusion injury^28^ and cardiac hypertrophy^29^, HDAC6 inhibition has shown great promise for potential future treatment options.

Targeting HDAC6 is of crucial interest, due to its therapeutic potential in various incurable diseases for which suitable therapies are currently lacking. Therefore, this study aimed to develop and characterize a novel, fluorescent HDAC6-specific probe that can be applied as a screening tool in living cells.

## Results

### Design and synthesis of MeFluHyA (Methyl Fluorescent Hydroxamic Acid)

Pan-HDAC inhibitors typically comprise three key structural elements: a zinc-chelating moiety targeting the metal ion in the active site of the enzyme, a linker and a bulky capping group interacting with the enzyme surface. Addition of a large, polycyclic moiety to the capping group, in combination with a phenyl hydroxamic acid pharmacophore, has been shown to enhance selectivity toward HDAC6 by targeting the enzyme’s shorter and wider catalytic pocket more effectively^30^. In order to generate a fluorescent probe targeting HDAC6 selectively, we replaced the bulky capping group of phenyl hydroxamic acid-based drugs with a fluorophore, Cy5 (**Figure 1A)**. This far-red fluorophore is cell-permeable, not toxic and allows live-cell imaging experiments with low excitation energy and deep penetration depth while facilitating orthogonality with eGFP-based colocalization studies.

**Figure 1:**
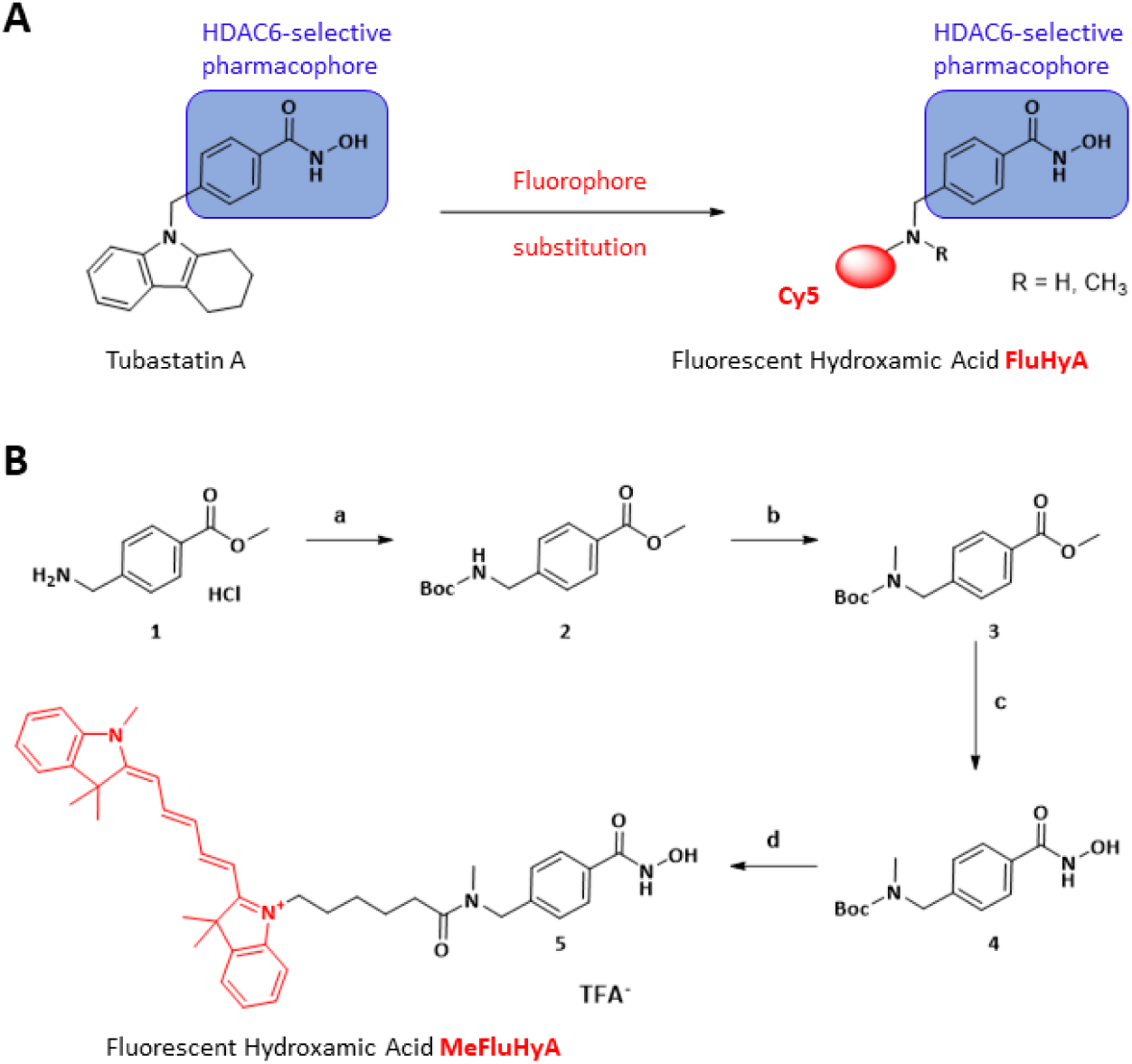
Strategy for fluorescent probe synthesis: **A)** General strategy for the synthesis of fluorescent hydroxamic acids by substituting the bulky capping group of an HDAC6-selective hydroxamic acid with a fluorophore (Cy5) and conserving the phenylhydroxamic acid group as pharmacophore to ensure HDAC6-selectivity. **B)** Chemical synthesis of MeFluHyA: a) Boc_2_O, THF/H2O, ACN (94%); b) NaH, CH_3_I, DMF, 0°C -> RT (79%); c) NH_2_OH, NaOH, THF/MeOH, 0°C -> RT (89%); d,e) 4 M HCl/1,4-dioxane, then NHS-Cy5-ester, TEA, DCM/DMSO (93%).

We synthesized a small library of four chemical probes, derived from commercially available methyl-4-aminomethyl benzoate hydrochloride, yielding Fluorescent Hydroxamic Acid (FluHyA) derivatives (see Supplementary Information and **SI Figure 1**). Following protection^31^, the nitrogen of compound **2** was methylated, the resulting compound **3** converted into a hydroxamic acid derivative and subsequently coupled to a Cy5 NHS-ester to afford the lead compound MeFluHyA (**Figure 1B**).

### Evaluation of MeFluHyA and its HDAC6-selectivity

To evaluate the potency and selectivity of MeFluHyA (and the other FluHyAs, see SI Fig. 2), we screened the inhibitory activity (IC_50_) in biochemical, fluorogenic *in vitro* assays against HDAC6 (**Figure 2A**) and other HDAC isoforms (**Figure 2B**, **SI Figure 2, SI Table 1**). The panel included 10 HDACs and sirtuin 2 (SIRT2), as this HDAC isoform can also deacetylate α-tubulin^32^. Within this set of HDACs, MeFluHyA demonstrated superior against HDAC6 as represented in the heatmap (**Figure 2B**). It was therefore selected as lead compound due to its biochemical and imaging (see below) properties. To investigate downstream effects of HDAC6-selective inhibition, we treated human SH-SY5Y neuroblastoma cells with MeFluHyA in a dose-dependent (**Figure 2C, 2F**) and time-dependent manner (**SI Figure 3A, 3B**). Increased acetylated α-tubulin revealed inhibition of HDAC6, while acetylated histone 3 levels showed non-selective HDAC class I inhibition. MeFluHyA increased α-tubulin acetylation at micromolar concentrations and exhibited minimal class I HDAC inhibition, markedly reduced at micromolar concentrations compared to Tubastatin A, a well-characterized and widely used HDAC6-selective inhibitor^30^ (Figure 1A). Notably, MeFluHyA and Tubastatin A treatment resulted in increased α-tubulin acetylation already after 30 min (**SI Figure 3A**). Taken together, these results indicate that the biological effect of HDAC6 inhibition is fast and selective, highlighting the utility of competitive probes for the identification of novel HDAC6 inhibitors.

**Figure 2:**
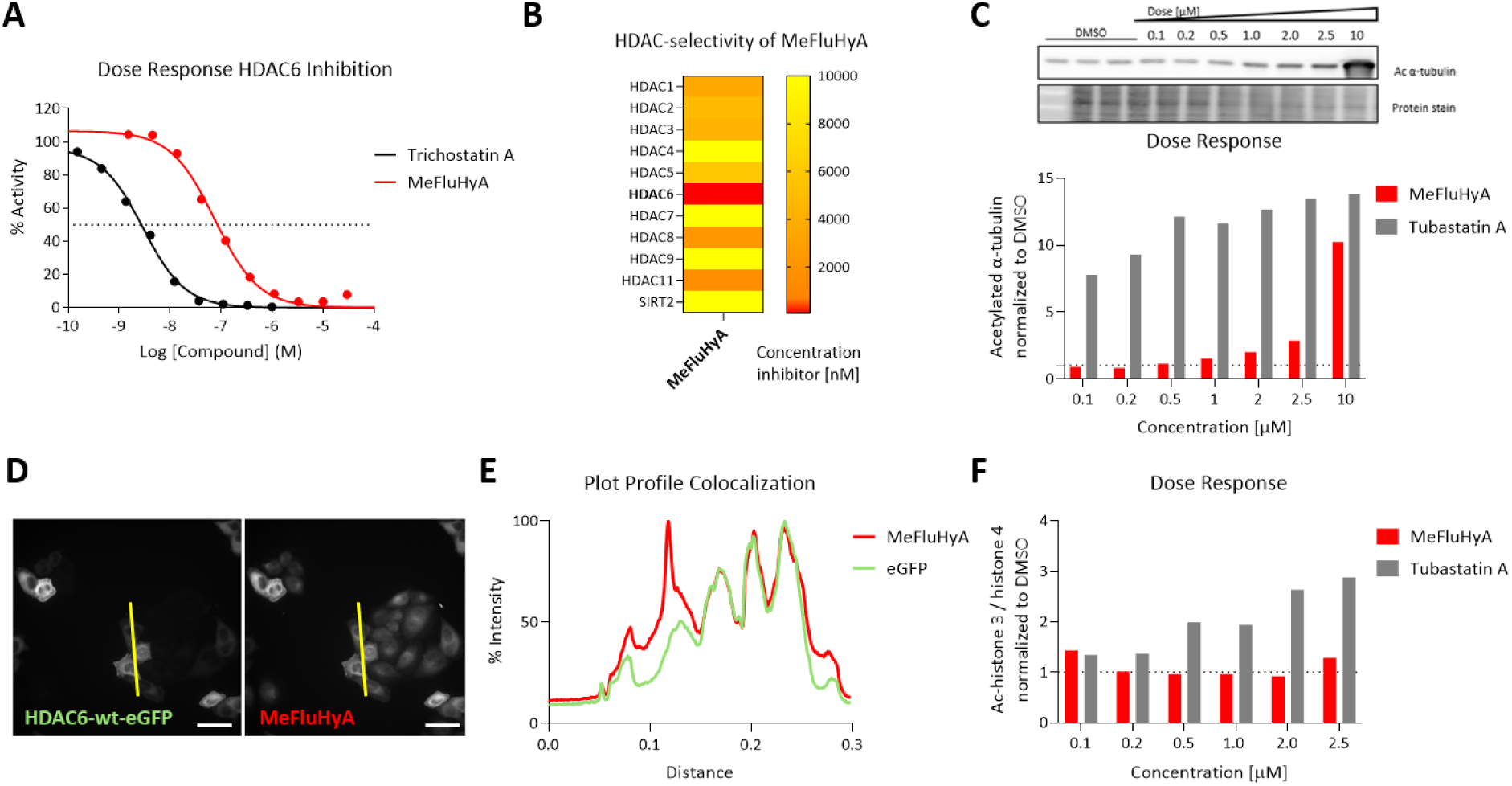
Characterization of MeFluHyA in an HDAC profiling *in vitro* assay, dose-dependent HDAC6 downstream inhibitory activity as well as fluorescent imaging properties in cells. **A)** MeFluHyA and Trichostatin A show dose-dependent HDAC6 inhibition with sigmoidal dose-response curve fitting in a biochemical HDAC6 assay. Curve fitting and IC_50_ values can be found in the supplementary material. **B)** Heatmap representing HDAC inhibition potency, with MeFluHyA most potently inhibiting HDAC6. **C)** Western blots illustrating the dose-dependent effect of MeFluHyA on acetylated α-tubulin as readout for HDAC6 inhibition after 6 h incubation in SH-SY5Y cells. **D)** HeLa cells were transfected using FuGene transfection reagent with a HDAC6-wt-eGFP plasmid for 24 h and incubated with the probe (50 nM) for 30 min, fixed and imaged on an Operetta CLS High Content microscope. Scale bar = 50 μm. **E)** Plot profile of eGFP (HDAC6) and Cy5 (MeFluHyA), the line in (D) shows the location of the ROI depicted in the plot profile. **F)** Quantifciation of the dose-dependent effect of MeFluHyA on acetylated histone 3 over histone 4 as readout for no apparent pan-HDAC inhibition after 6 h incubation in SH-SY5Y cells.

**Figure 3:**
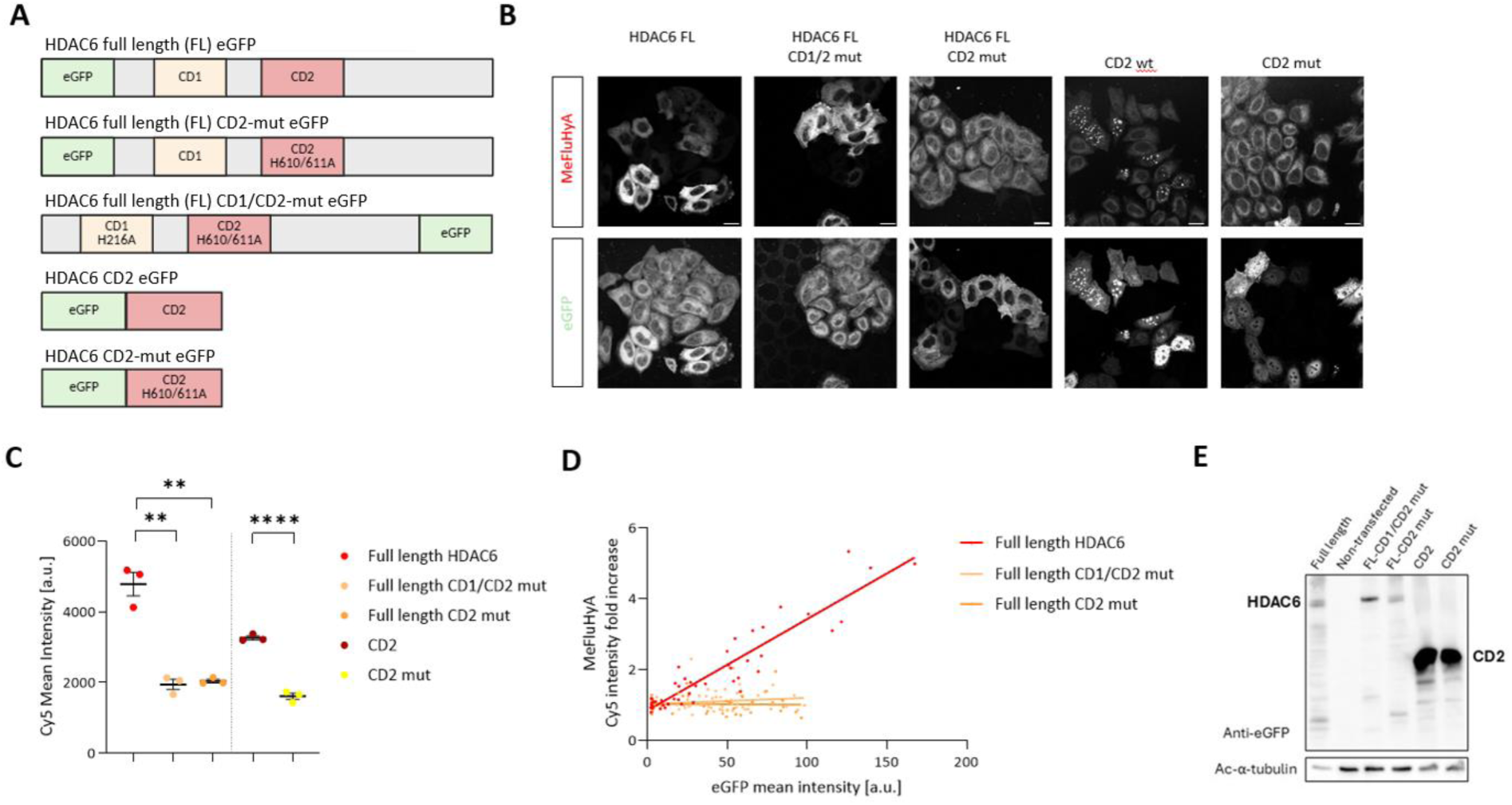
The second catalytic domain (CD2) of HDAC6 is the binding site of the fluorescent probes. **A)** Schematic representation of the HDAC6 plasmids used. HDAC6-full length-eGFP: HDAC6-full length CD2 mutated (H610/611A)–eGFP; HDAC6-full length CD1/ CD2 mutated (H216A and H610/611A)–eGFP; HDAC6-CD2-eGFP and HDAC6-CD2 mutated (H610/611A)-eGFP. **B)** Treatment of HeLa cells transiently overexpressing one of the HDAC6-plasmids with 200 nM MeFluHyA using FuGene transfection reagent Left: HDAC6-wt-eGFP, middle: HDAC6-CD1/CD2-mut-eGFP and right: HDAC6-CD2-mut-eGFP mutant. Figures indicate a correlation between HDAC6 and the probe but an absence of correlation between CD2 and CD1/CD2 mutated HDAC6. Scale bar = 20 µM, images Leica SP8 confocal microscope (63x magnification, n = 3 biological replicates). **C)** Correlation between different plasmids and MeFluHyA indicating binding depending on functionality of CD2 (Unpaired t-test, n = 3 biological replicates, ** p<0.01, **** p<0.0001). **D)** Cy5 mean intensity of MeFluHyA in HeLa cells overexpressing different HDAC6 plasmids. **E)** Western blot of HeLa protein extract after 24 h transfection of the different constructs and non-transfected control, probed with antibodies against eGFP and acetylated α-tubulin, highlighting successful transfection of the constructs and functional effect of HDAC6 overexpression on acetylated α-tubulin. All data are presented as mean ± SEM.

Next, we evaluated the imaging properties of MeFluHyA to further confirm HDAC6-selectivity. HeLa cells were transiently overexpressed with full-length wild-type HDAC6 tagged with eGFP, and signal intensities of MeFluHyA were compared between transfected and non-transfected cells across a concentration range (100 nM to 1 µM). For all concentrations, MeFluHyA exhibited a stronger fluorescence signal in transfected cells relative to non-transfected cells (**SI Figure 4**). The strongest correlation between MeFluHyA and HDAC6-eGFP fluorescence intensities was observed at the lowest concentration (100 nM). Furthermore, fluorescence intensity plot profiles showed a clear colocalization pattern between MeFluHyA and HDAC6-eGFP (**Figure 2D, E**). Additionally, MeFluHyA colocalizes with a commercial HDAC6 antibody (**SI Figure 5**).

**Figure 4:**
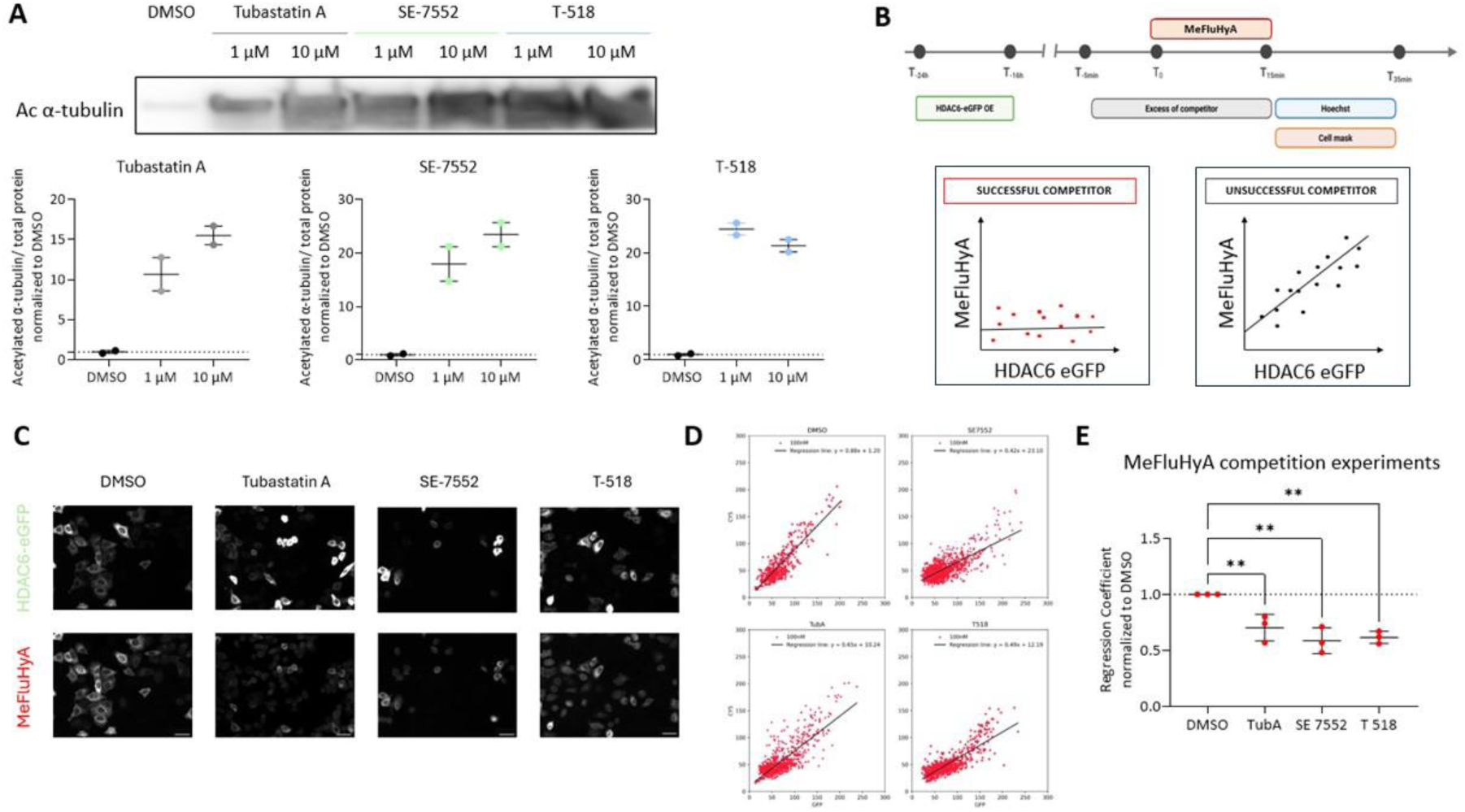
Fluorescent probes are able to identify HDAC6-targetting drugs in HDAC6-overexpressing HeLa cells. **A)** Dose-response curves of three different HDAC6 inhibitors (Tubastatin A, SE 7552 and T 518) using acetyl-α-tubulin as functional read-out. **B)** Experimental paradigm of competition experiments using HeLa cells transiently overexpressing full length HDAC6-eGFP (using Lipofectamine 3000 transfection reagent). **C)** Treatment of HeLa cells transiently overexpressing HDAC6-full length-eGFP with 100 nM MeFluHyA according to experimental paradigm A using Tubastatin A, SE 7552 or T-518 as competitor. Images were taken on a Zeiss cell discoverer 7 microscope (20x magnification, scale bar= 10 µm) **D and E**) Correlation plots and quantification of competition experiments. The correlation between HDAC6-full length-eGFP and MeFluHyA is decreased upon competition with excess of an HDAC6-inhibitor (unpaired t-test, ** p<0.01, n = 3 biological replicates). All data are presented as mean ± SEM.

### Investigation of MeFluHyA binding properties

To investigate which catalytic domain (CD1 or CD2) MeFluHyA targets, we transiently overexpressed different constructs of HDAC6-eGFP (**Figure 3A**). Mutations in the previously used HDAC6-full length plasmid were introduced in CD2 (H610/611A) and CD1 (H216A) to inactivate their catalytic activity. Furthermore, we included CD2 excerpts, both wild type and mutated (H610/611A).

Consistent with the results shown in Figure 2, fluorescent intensities of HDAC6-eGFP and MeFluHyA were positively correlated. Interestingly, this correlation was lost upon catalytic inactivation of CD2, indicating specific binding of the fluorescent probe to catalytically active CD2 (**Figure 3 B, C, D**). To validate successful expression of the constructs, HeLa cell lysates were probed for eGFP and acetylated α-tubulin. In line with the catalytic inactivation, transfection of the mutated plasmids did not influence α-tubulin acetylation levels (**Figure 3E**). This is in line with previous findings, that most HDAC6 inhibitors target CD2, while CD1 exhibits much narrower specificity^33^.

### MeFluHyA as a tool to identify successful HDAC6 inhibitors

As MeFluHyA targets the second catalytic domain of HDAC6, responsible for the deacetylation of α-tubulin, we aimed to evaluate its potential as a fluorescent probe to identify successful HDAC6 inhibitors in a competitive assay. For this, we used three commercially available HDAC6 inhibitors, Tubastatin A, SE 7552 and T 518. All demonstrated increased acetylation of α-tubulin on western blot (**Figure 4A**). All three inhibitors were able to occupy the catalytic site of HDAC6, resulting in decreased binding of MeFluHyA (**Figure 4B**) despite their differences in chemical structure^16^ (**Figure 4C, SI figure 6**). To quantify this effect, we analyzed the correlation between HDAC6-eGFP and MeFluHyA (Cy5) fluorescence signals. A decrease in the regression coefficient of this correlation served as an indicator of successful HDAC6 inhibition, allowing us to identify compounds that effectively compete with MeFluHyA for binding to HDAC6 (**Figure 4D-E**).

## Discussion

In this study, we present MeFluHyA, a novel fluorescent probe specifically engineered to enable high-throughput identification of HDAC6 inhibitors in cells. By integrating a Cy5 fluorophore into a phenyl hydroxamic acid scaffold, we designed a probe that not only retains HDAC6 selectivity but also facilitates high-throughput identification of HDAC6 inhibitors. While fluorophore-conjugated hydroxamic acid derivatives have previously been explored, many suffer from fundamental limitations that restrict their utility in dynamic cellular assays due to their excitation at a low wavelength, and therefore causing tissue damage and low penetration depth^34–37^. Even though Meng et al. introduced Cy5.5-labeled derivatives such as LBH589-Cy5.5, which demonstrated potent HDAC inhibition in nuclear extracts (IC₅₀ = 9.6 nM)^38^, the presence of three anionic sulfo groups limits its cell-permeability. Thus, it is unsuitable to be used in live-cell competition assays and restricts its applicability to fixed-cell or lysate-based formats. In another, recent study by Nguyen and colleagues (2026) the authors developed a rhodamine fluorescent probe based on Nexturastat, called 6SiR-C3-NextA, that successfully labeled HDAC6 in multiple cell lines, however, only showed partial overlap when co-stained with an HDAC6 antibody, suggesting that further probe refinement could enhance the probe’s features ^39^.

To overcome previous limitations, we designed MeFluHyA with a Cy5 fluorophore, which offers both cell permeability and compatibility with far-red imaging and a quick labeling procedure. This allows orthogonal compatibility with commonly used fluorescent tags such as eGFP, increasing the probe’s applicability in multiplex imaging experiments and potentially application to different species.

Our biochemical assays confirmed that MeFluHyA selectively targets HDAC6 over other HDAC isoforms, including class I HDACs and SIRT2. In addition, the probe resulted in increased α-tubulin acetylation, a functional readout of HDAC6 inhibition without significantly affecting histone H3 acetylation, indicating minimal off-target activity. These findings validate the probe’s selectivity. Different imaging experiments further demonstrated that MeFluHyA colocalizes with HDAC6-eGFP and a commercial HDAC6 antibody, confirming its cellular selectivity. Domain-specific mutagenesis experiments revealed that MeFluHyA binds selectively to the catalytically active CD2 domain of HDAC6, which is responsible for α-tubulin deacetylation.

Importantly, MeFluHyA binds HDAC6 at low concentrations without significantly inhibiting its activity, which is critical for its intended use. Unlike traditional inhibitors, our probe is not aimed to impair HDAC6 function, but rather to serve as a competitive readout for identifying compounds that do, as probes that inhibit HDAC6 can confound screening results by masking the effects of candidate inhibitors. This property, combined with its domain-specific binding to the catalytically active CD2 domain, positions MeFluHyA as a selective tool for screening platforms.

Overall, MeFluHyA addresses key shortcomings of previous probes and introduces a new strategy for competitive inhibitor screening in a cellular model. MeFluHyA demonstrated effective identification of different known commercially available HDAC6 inhibitors in our competition assay, highlighting its potential as a screening tool. Future work should explore its integration into automated imaging pipelines and its performance across diverse cellular models to further validate its utility in drug discovery.

In summary, MeFluHyA is a selective, cell-permeable fluorescent probe that facilitates the identification of novel HDAC6 inhibitors. Its application in competitive screening platforms provides a future avenue for discovery processes to examine compound potency and selectivity of novel HDAC6 inhibitors, hopefully facilitating drug discovery efforts to find treatments for neurodegenerative diseases.

## Materials and methods

### Chemistry

Intermediate compounds were analyzed using TLC, ^1^H- and ^13^C-NMR, analytical HPLC, LC/MS and/or HRMS. Final product formation was confirmed using HRMS and analytical HPLC. Unless specified otherwise, ^1^H NMR spectra were recorded on Bruker Avance III HD 400 spectrometer (working at 400 MHz, console with a Bruker Ascend™ 400 magnet, equipped with a 5 mm PABBO BB/19F-1H/D probe with z-gradients and ATM accessory for Automatic Tuning and Matching). Samples were dissolved in CDCl_3_, CD_3_OD or DMSO-*_d6_* and tetramethylsilane was used as an internal standard. The *δ*-values are expressed in ppm. The following abbreviations are used in reporting NMR data: *s* (singlet), *d* (doublet), *t* (triplet), *q* (quartet), *m* (multiplet), and as combinations thereof, with “*br*” indicating a broadened peak(s) and “*app*” indicating apparent. ^13^C-NMR spectra were recorded on Bruker Avance III HD 400 (working at 100 MHz) spectrometer, unless specified otherwise. The deuterated solvents were used as internal standard (CDCl_3_: 77.16 ppm, t; CD_3_OD: 49.01 ppm, septet; DMSO-*_d6_*: 39.52 ppm, septet). The δ-values are expressed in ppm. HR-MS spectra were acquired on a quadrupole orthogonal acceleration time-of-flight mass spectrometer (Synapt G2 HDMS, Waters, Milford, MA). Samples were dissolved in a mixture of 1:1 (v/v) acetonitrile:water and infused at 5 μL/min. Spectra were obtained in positive or negative ionization mode with a resolution of 15000 (fwhm) using leucine enkephalin as lock mass. HPLC: Unless specified otherwise, the purification of the final compounds was performed using a Buchï C-850 FlashPrep equipped with a Robusta^®^ C18 (5µ-150mm x 21.2 mmID) prep HPLC column. Conditions are specified in the individual experiments. Materials: All reagents were obtained from commercially available sources and were used as purchased without further purification. All moisture-sensitive reactions were carried out under nitrogen atmosphere and in oven-dried glassware. Yields refer to isolated compounds after quantitative conversion or after chromatography. Cy5 NHS-Ester was purchased from Lumiprobe. Detailed synthesis can be found in supplementary materials.

### *In vitro* compound screening with recombinant HDAC proteins

Experiments were carried out by Reaction Biology Corporation (Woodbridge, CT, USA). Briefly, recombinant protein, fluorogenic peptide and respective compounds were incubated at different concentrations. Deacetylase activity was assessed by measuring fluorophore release and inhibition resulted in decreased fluorescence. A counter screen was performed to exclude cross-reactivity of the fluorogenic FluHyA compounds with the assay readout. HDAC6 screening has been performed at 10 dose singlet screen, starting from 30 µM compound with a 3-fold dilution factor. All other targets have been screened at a 5-dose singlet application, starting from a 10 µM concentration with a 10-fold dilution factor. HDAC reference compounds Trichostatin A (TSA) and TMP269 were tested in 10-dose IC50 mode with 3-fold serial dilution starting at either 1 or 10 µM.

### Cell culture

SH-SY5Y neuroblastoma cells (ATCC #: CRL-2266) were cultivated at 37°, 5% CO_2_ in Dulbecco’s Minimal Essential Medium and F12 medium (DMEM/F-12 GlutaMax, Gibco^TM^ #: 31331), supplemented with 10% fetal bovine serum (FBS, Gibco^TM^ #: 10270098), 1% Pen/Strep (Gibco^TM^ #: 15070063), 1% MEM Non-Essential Amino Acids (MEM NEAA, Gibco^TM^ #: 11140050). HeLa cells (abcam #: ab255448) were cultivated at 37°, 5% CO_2_ in DMEM, high glucose with GlutaMAX^TM^ (Gibco^TM^ #: 10566016), with 10% FBS and 1% Pen/Strep. Cells were routinely passaged with 0.05% trypsin-EDTA (Gibco^TM^ #: 25300054). Cell cultures were subjected to monthly mycoplasma testing.

Transfection was performed using FuGene® HD Transfection Reagent (Promega, #: E2311) or Lipofectamine^TM^ 3000 Transfection Reagent (Invitrogen^TM^ #: L3000001), according to the manufacturer’s guidelines.

### Compound treatment

The fluorescent compound was diluted from a 1 mM stock concentration in DMSO. Tubastatin A was purchased from Selleck Chemicals (#S8049), SE-7552 from MedChemExpress (HY-161305) and T-518 from MedChemExpress (HY-161307). Vehicle-treated cells were exposed to the same concentration of DMSO.

### Competition experiments

HeLa cells were transfected using Lipofectamine^TM^ 3000 Transfection Reagent (Invitrogen^TM^ L3000001), according to manufacturer’s guidelines. 24 hours later, an excess of competitors was applied for 5 min. Next, the MeFluHyA (100 nM) was added for 15 min. After 20 min incubation with Hoechst (50 nM, Thermo Scientific™, 62249) and Cell mask Orange (1:10 000, Invitrogen^TM^, #: C10045) in Dulbecco’s phosphate buffered saline (DPBS), cells were fixed in 4% PFA. Images were acquired on a Zeiss Cell Discoverer 7 microscope at 20x magnification.

Image analysis was performed using ImageJ. For this, cells were masked using the cell mask signal, and regions of interest were determined based on Hoechst signal. Then, intensities of HDAC6 (eGFP) and MeFluHyA (Cy5) were measured in each cell individually, and correlated.

### Protein extraction and Western blot

After treatment, cells were washed twice with DPBS on ice, and homogenized using EpiQick Total Histone Extraction Kit (EpigenTek # OP-0006-100) to extract the cytosolic and histone fraction, according to the manufacturer’s protocol. Briefly, pre-lysis buffer (supplemented with cOmplete™ Mini, EDTA-free Protease Inhibitor Cocktail from Roche, Merck #11836153001, and PhosStop™ Phosphatase Inhibitor from Roche, Merck #: 4906845001) was added and incubated for 5 min, cells were scraped and transferred into a microcentrifuge tube and incubated for 30 min on ice. The histone fraction was pelleted by centrifugation at 12.000 rpm for 5 min at 4 °C. The supernatant was kept as the cytosolic fraction and protein concentration was determined using Thermo Scientific™ Pierce™ BCA Protein Assay Kit (#23225). Samples were resuspended in 5X Pierce™ Lane Marker Reducing Sample Buffer and boiled at 95 °C for 10 min. 4-20% Criterion™ TGX™ Precast Mini protein gels (BioRad, #4568096) were loaded with 5 μg protein, and separated at 120 V for 50 min. Transfer was performed using Trans-Blot Turbo System (BioRad) using Trans-Blot Turbo Mini 0.2 µm PVDF Transfer Packs (Biorad, #1704156) with a 30 min transfer. After blocking the membrane for 1 h in 5% BSA in Tris-buffered saline solution with 0.1% Tween 20 (TBS-T), incubation of primary antibody (in 1% BSA and TBS-T) was performed for 1 to 2h at room temperature or overnight at 4 °C. After 3x 5 min TBS-T washes, the membrane was incubated with complementary secondary antibody (Dako, P0447 and P0448, 1:2000 dilution) for 1 h at room temperature, followed by two TBS-T and one TBS wash. Blots were visualized using ECL SuperSignal (ThermoFisher, #32106) and an ImageQuant LAS 4000 imager (GE Healthcare). Antibodies used were: Anti-acetyl α-tubulin (Sigma, T6793; 1:5000), Anti-β-Actin (Sigma, A5441, 1:1000), Anti-acetyl-Histone H3 (Lys9) (Millipore, 06-942, 1:1000), Anti-Histone H4 (abcam, ab10158, 1:1000), Anti-eGFP (CST, 2956S, 1:1000).

### Immunocytochemistry

Cells were fixed in 4% PFA for 15 min at room temperature. After blocking in 5% normal donkey serum (NDS; Sigma) in 0.1% Triton X-100 in PBS (PBS-T) for 1 h, primary HDAC6 antibody (abcam, ab133493 1:1000) was incubated overnight at 4 °C in 1% NDS in PBS-T, followed by 3× 10 min DPBS washes and secondary antibody treatment with Alexa Fluor 488/555 antibodies (Invitrogen, 1:1000) for 2h at 37 °C. Next, NucBlue^TM^ Live ReadyProbes^TM^ Reagent (Hoechst 33342) (Invitrogen^TM^, #: R37605) was used 10 min. For mounting, ProLong Diamond Antifade Mountant was used (Invitrogen #: P36962) and the mounted coverslips were kept for 24h for equilibration until imaging. All images were acquired on a Leica SPM8 confocal microscope or on a Perkin Elmer Operetta (Perkin Elmer).

### Statistics

All statistical analyses were performed using GraphPad Prism software version 9 (GraphPad software Inc., California, USA). Comparisons between two groups were conducted using an unpaired, two-tailed Student’s *t*-test, while one-way ANOVA was used for multiple group comparisons. Data are presented as mean ± SEM. Statistical significance was set at: * p < 0.05, ** p < 0.01, *** p < 0.001, **** p < 0.0001.

## Acknowledgements

We would like to thank Nicole Hersmus, Wendy Vermeire, Thibaut Burg, Alessio Silva, Thomas Moens and Frederik Rombouts for additional help and discussions.

## Author Contributions

Y.E.K., M.S. and L.V.D.B. conceptualized the study. Y.E.K. performed chemical and biological experiments, imaging and wrote the manuscript. J.G. performed biological experiments, image acquisition and processing and wrote the manuscript. A.S. helped performing biological and imaging experiments and automated analysis. C.B., and E.I. helped with compound synthesis. J.V.L. helped performing biological experiments, image processing and statistical analysis. R.P. helped performing biological experiments. W.D.B., S.V. contributed in synthetical planning and discussions. P.V.D., J.M.H. and M.C. contributed to discussions. M.C. provided HDAC6-GFP plasmids. All authors provided feedback to the manuscript, read and agreed to the final version of this manuscript.

## Funding Sources

Y.E.K. was a FWO PhD-fellow SB (#1S50320N), J. G. is a FWO PhD-fellow Fundamental Research (#1124325N), C. B. was a FWO PhD-fellow SB (#1SB6521N), E.I. was a FWO Post-Doc fellow (#12Z6620N), R.P. was a FWO PhD-fellow SB (#1S59317N). This research was supported by VIB, KU Leuven (C1 and “Opening the Future” Fund), the “Fund for Scientific Research Flanders” (FWO-Vlaanderen, G026924N and G088523N), the Thierry Latran Foundation, the “Association Belge contre les Maladies neuro-Musculaires – aide à la recherché ASBL” (ABMM), the Muscular Dystrophy Association (MDA), the ALS Liga België (A Cure for ALS), Target ALS and the ALS Association (ALSA). PVD holds a fundamental clinical investigatorship of KU Leuven. PVD and LVDB are supported by the Dr. Helen Camerlynck Chair, the ALS Liga België and the KU Leuven funds “Een Hart voor ALS” and the “Laeversfonds voor ALS Onderzoek”.

## Competing Interests Statement

L.V.D.B. is head of the Scientific Advisory Board of Augustine Therapeutics (Leuven, Belgium) developing improved HDAC6 inhibitors for peripheral neuropathies. Augustine Therapeutics was not involved in any aspect of this study. L.V.D.B is also part of the Investment Advisory Board of Droia Ventures (Meise, Belgium). PVD has served in advisory boards for Biogen, CSL Behring, Alexion Pharmaceuticals, Ferrer, QurAlis, Cytokinetics, argenx, UCB, Muna Therapeutics, Alector, Augustine Therapeutics, VectorY, Zambon, Amylyx, Novartis, Prilenia, Verge Genomics, Sapreme Technologies, Trace Neuroscience, NRG Therapeutics (paid to institution). PVD has received speaker fees from Biogen and Amylyx (paid to institution).

---

**Supplementary Figure 1:**
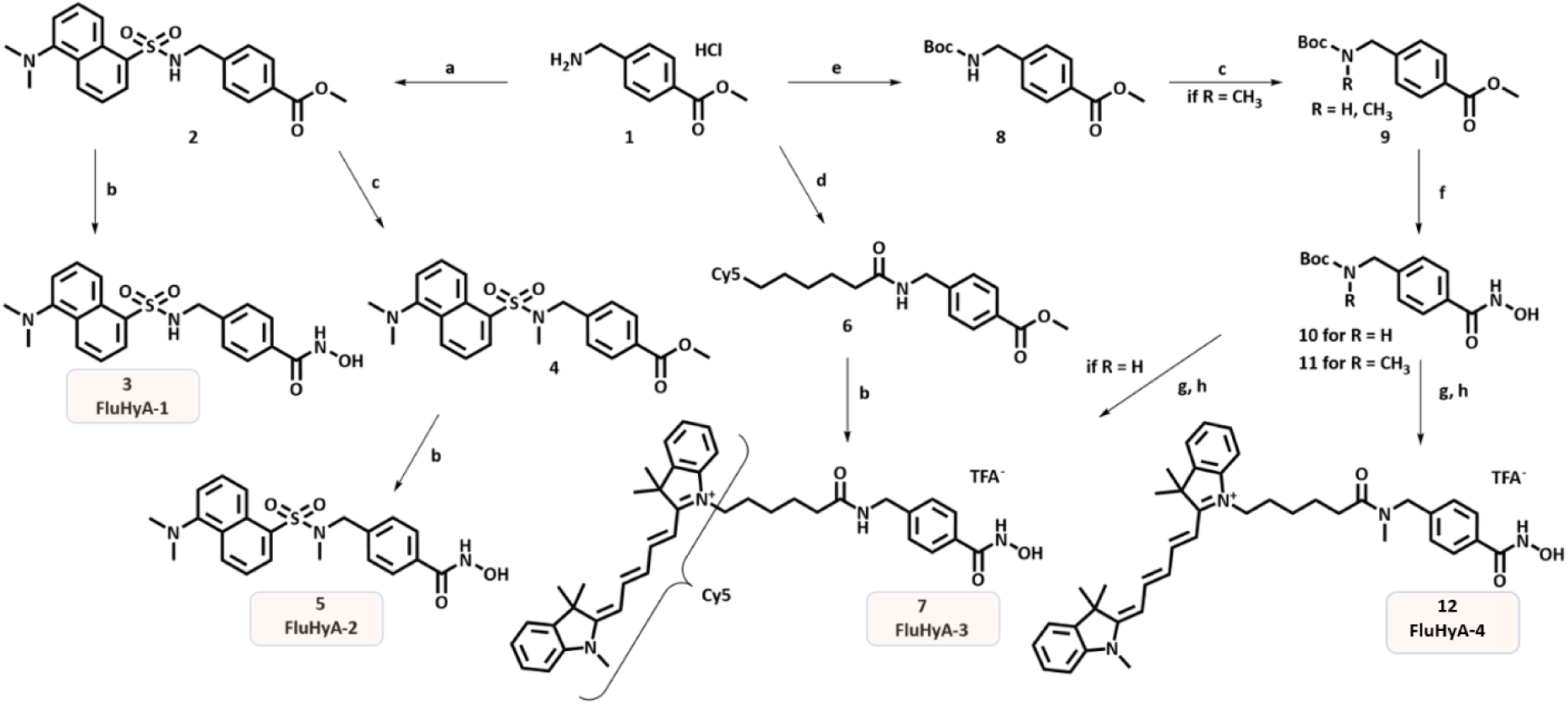
Chemical synthesis to obtain fluorescent hydroxamic acid-based compounds FluHyA 1 through 4. **a)** Dansylchloride, TEA, THF, 0°C -> RT (89%); **b)** H_2_NOH, NaOH, THF/MeOH (61% for 3, 57% for 5, 11% for 7); **c)** NaH, CH_3_I, DMF, 0°C -> RT (23% for 4; 79% for 9); **d)** Cy5-NHS-ester, THF, 0°C -> RT (quant.); **e)** Boc_2_O, THF/H_2_O, ACN (94%); **f)** NH_2_OH, NaOH, THF/MeOH, 0°C -> RT (89%); **g,h)** for R = H HCl (gaseous) generated *in situ*, then NHS-Cy5-ester, TEA, DCM (96%); for R = CH_3_ 4 M HCl/1,4-dioxane, then NHS-Cy5-ester, TEA, DCM/DMSO (93%).

**Supplementary Figure 2:**
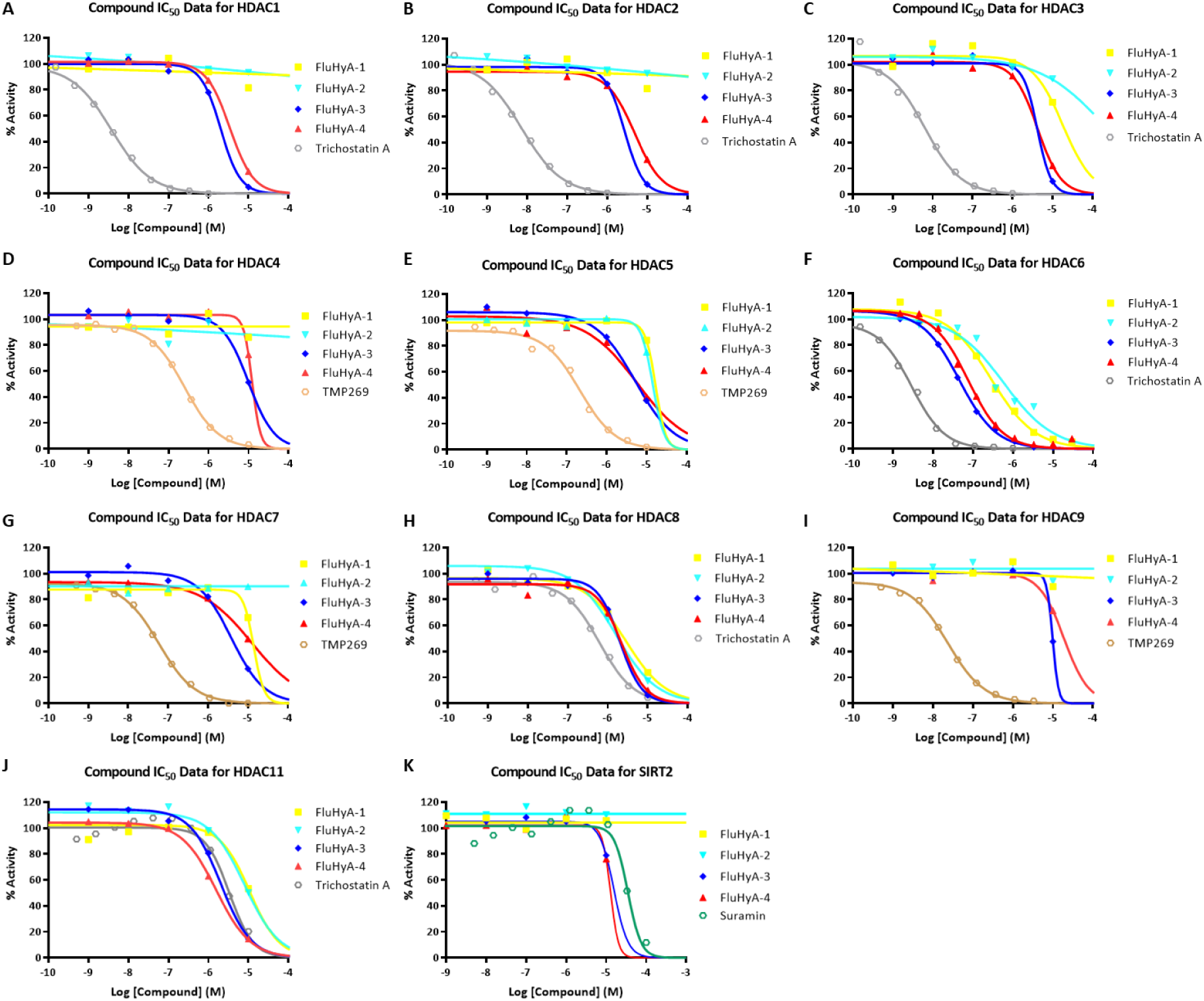
HDAC profiling IC_50_ assay. Compound IC_50_ values and reference compounds. HDAC inhibition fluorescence assays were performed by Reaction Biology Corporation. Compounds were tested in a singlicate 10-dose IC_50_ mode with 3-fold serial dilution starting from 30 µM for HDAC6; all other HDACs were tested in a singlicate 5-dose IC_50_ mode with 10-fold serial dilution starting from 10 µM.

**Supplementary Table 1:** IC_50_ values of MeFluHyA and comparison with control compounds.

| Target | Compound IC<sub>50</sub> (μM) | Control IC<sub>50</sub> (μM) |  |
| --- | --- | --- | --- |
|  | MeFluHyA | Control ID | IC<sub>50</sub> (μM) |
| HDAC1 | 3.43 | Trichostatin A | 0.004 |
| HDAC2 | 4.88 | Trichostatin A | 0.007 |
| HDAC3 | 4.21 | Trichostatin A | 0.007 |
| HDAC4 | >10 | TMP269 | 0.252 |
| HDAC5 | 5.82 | TMP269 | 0.214 |
| **HDAC6** | **0.075** | **Trichostatin A** | **0.003** |
| HDAC7 | >10 | TMP269 | 0.059 |
| HDAC8 | 2.24 | Trichostatin A | 0.621 |
| HDAC9 | >10 | TMP269 | 0.024 |
| HDAC11 | 1.57 | Trichostatin A | 2.32 |
| SIRT2 | >10 | Suramin | 34.9 |

**Supplementary Figure 3:**
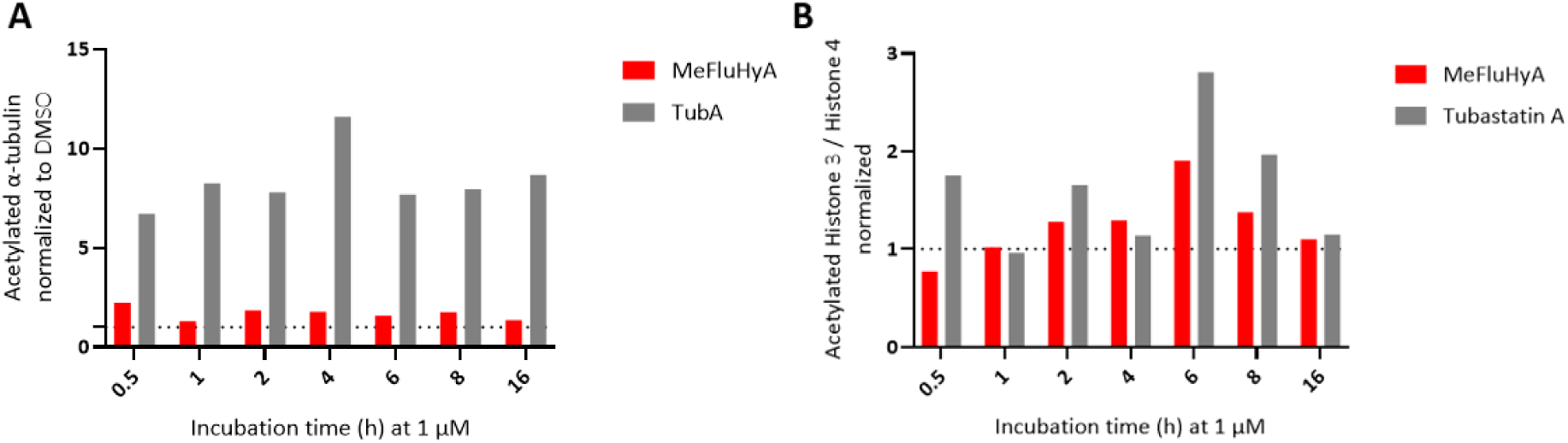
HDAC inhibition by MeFluHyA in SH-SY5Y cells. A) Time-response quantification of western blot of acetylated α-tubulin after HDAC6 inhibition by 1 µM of MeFluHyA in SH-SY5Y cells. **B)** Time-response quantification of western blot of histone-acetylation as a consequence of non-selective class I HDAC inhibition in SH-SY5Y cells.

**Supplementary Figure 4:**
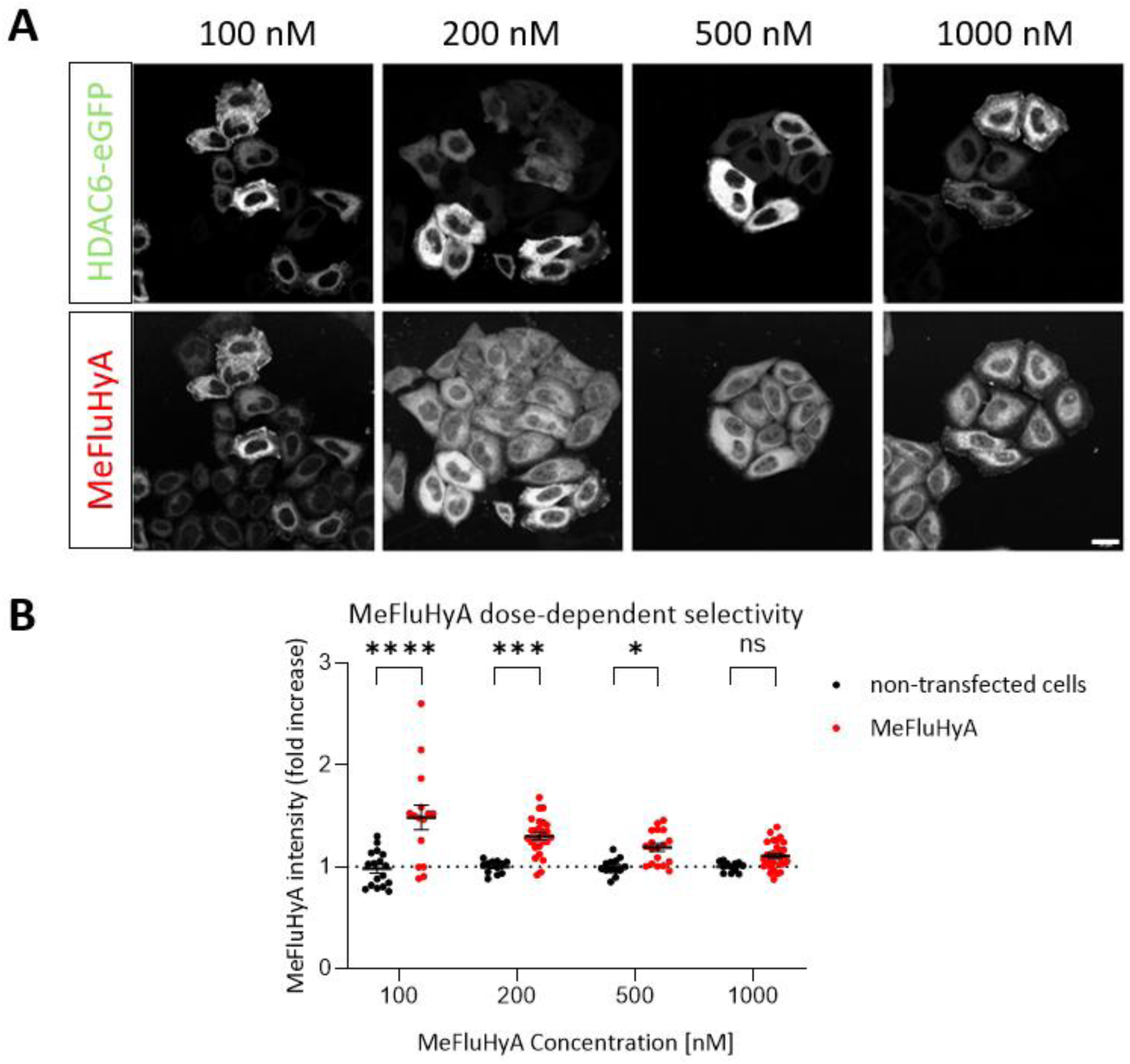
Colocalization of MeFluHyA with HDAC6-wt-eGFP. **A)** Fluorescent signal in HeLa cells after incubation with different concentration of MeFluHyA after 24h transient transfection with HDAC6-wt-eGFP using FuGene transfection reagent. Images were taken at a Leica SP8 confocal microscope. Scale bar = 20 µm. n = 12-29 cells; **B)** Quantification of the intensity signal in non-transfected and transfected (HDAC6-full length-eGFP) cells ranging from 100 nM to 1 µM MeFluHyA. Upon increased concentrations of MeFluHyA, the fold-change of Cy5 signal between HDAC6-eGFP and MeFluHyA is decreased, indicating decreased specific binding.

**Supplementary Figure 5:**
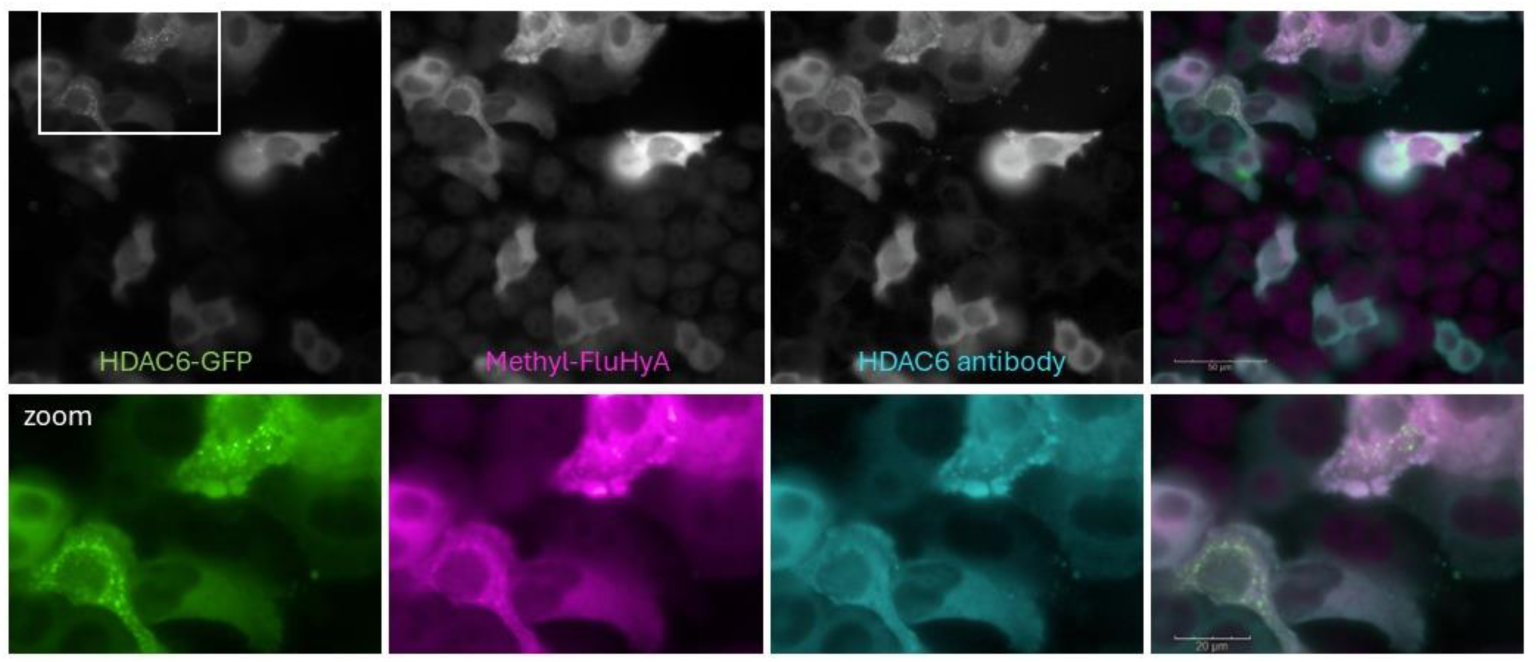
MeFluHyA colocalizes with HDAC6-eGFP and a commercial HDAC6 antibody. HeLa cells were transfected with HDAC6-full length-eGFP. The eGFP-signal colocalizes with the fluorescent probe, MeFluHyA and a commercial HDAC6 antibody (abcam, ab133493). Upper scale bar = 50 µm, lower scale bar = 20 µm.

**Supplementary Figure 6:**
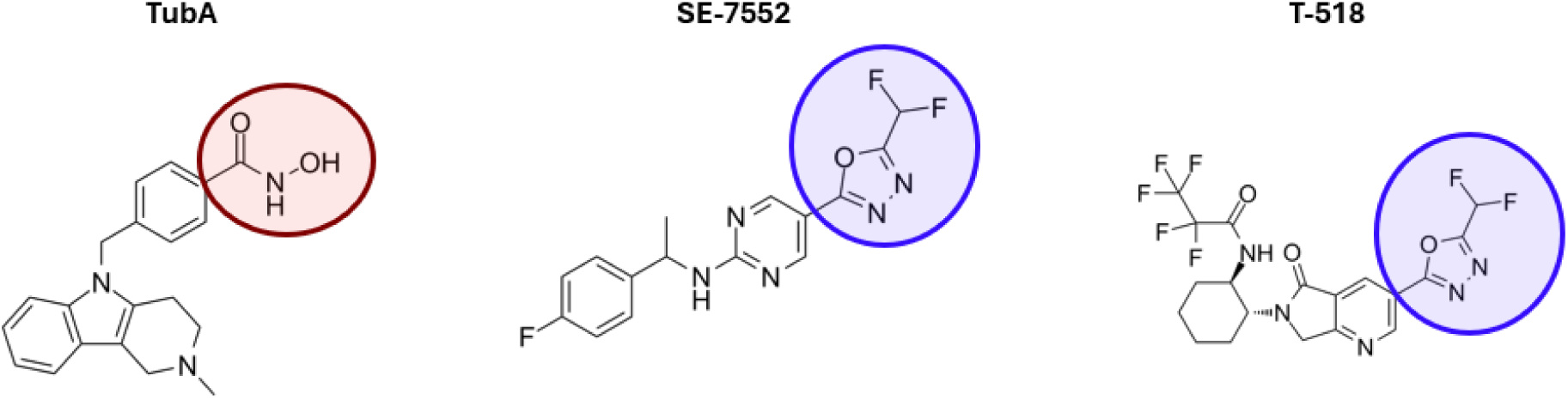
Chemical structures of commercially available HDAC6 inhibitors: Tubastatin A targets CD2 by its hydroxamic acid group (red). In contrast, SE 7552 and T 518 target via their fluoromethyl oxadiazole group (blue).

## References

1. Pulya, S. et al. Pharmacol Res 163, 105274 (2021).

2. Rossaert, E. & Van Den Bosch, L. Brain Res 1733, 146692 (2020).

3. Kawaguchi, Y. et al. Cell 115, 727–738 (2003).

4. Hubbert, C. et al. Nature 417, 455–458 (2002).

5. Noack, M. et al. Glia 62, 535–547 (2014).

6. Zhang, X. et al. Mol Cell 27, 197–213 (2007).

7. Parmigiani, R. B. et al. Proc Natl Acad Sci U S A 105, 9633–9638 (2008).

8. Kalinski, A. L. et al. J Cell Biol 218, 1871–1890 (2019).

9. Nakka, K. K. et al. Proc Natl Acad Sci U S A 112, E3374–3383 (2015).

10. Ding, H. et al. J Neurochem 106, 2119–2130 (2008).

11. Simoes-Pires, C. et al. Mol Neurodegener 8, 7 (2013).

12. Li, Y. et al. Eur J Med Chem 226, 113874 (2021).

13. Klingl, Y. E. et al. Br J Pharmacol 178, 1353–1372 (2021).

14. Benoy, V. et al. Brain 141, 673–687 (2018).

15. Prior, R. et al. Expert Opin Ther Targets 22, 993–1007 (2018).

16. van Eyll, J. et al. Expert Opin Ther Targets 28, 719–737 (2024).

17. Govindarajan, N. et al. EMBO Mol Med 5, 52–63 (2013).

18. Guedes-Dias, P. et al. Biochim Biophys Acta 1852, 2484–2493 (2015).

19. Lee, J. Y. et al. J Cell Biol 189, 671–679 (2010).

20. Van Helleputte, L. et al. Neurobiol Dis 111, 59–69 (2018)

21. d’Ydewalle, C., et al. Nat Med 17, 968–974 (2011).

22. Guo, W. et al. Nat Commun 8, 861 (2017).

23. Fazal, R. et al. EMBO J 40, e106177 (2021).

24. Stoklund Dittlau, K., et al. Stem Cell Reports 16, 2213–2227 (2021).

25. Tzeplaeff, L. et al. Cells 12 (2023).

26. Aldana-Masangkay, G. I. & Sakamoto, K. M. J Biomed Biotechnol 2011, 875824 (2011).

27. Ranjbarvaziri, S. et al. Nat Commun 15, 1352 (2024).

28. Baumgardt, S. L. et al. Cardiovasc Res 120, 1456–1471 (2024).

29. Yao, J. et al. J Pathol 266, 217–229 (2025).

30. Butler, K. V. & Kozikowski, A. P. Curr Pharm Des 14, 505–528 (2008).

31. Verschueren, R. H. et al. Org Biomol Chem 19, 5782–5787 (2021).

32. Skoge, R. H. et al. DNA Repair (Amst) 23, 33–38 (2014).

33. Osko, J. D. & Christianson, D. W. Biochemistry 58, 4912–4924 (2019).

34. Padilla-Coley, S. et al. Bioorg Med Chem Lett 47, 128207 (2021).

35. Zhang, Y. et al. Eur J Med Chem 141, 596–602 (2017).

36. Li, M. et al. Chem Commun (Camb) 58, 1938–1941 (2022).

37. Raudszus, R. et al. Bioorg Med Chem 27, 115039 (2019).

38. Meng, Q. et al. Mol Pharm 12, 2469–2476 (2015).

39. Nguyen, et al., J. Am. Chem. Soc. 148, 24, 24637–24645 (2026).

